# Fluorescence photography of patterns and waves of bacterial adaptation at high antibiotic doses

**DOI:** 10.1101/806232

**Authors:** Carlos Reding, Mark Hewlett, Tobias Bergmiller, Ivana Gudelj, Robert Beardmore

**Affiliations:** Biosciences, College of Life and Environmental Sciences, University of Exeter, Exeter EX4 4QD, UK

## Abstract

Fisher suggested advantageous genes would spread through populations as a wave so we sought genetic waves in evolving populations, as follows. By fusing a fluorescent marker to a drug efflux protein (AcrB) whose expression provides *Escherichia coli* with resistance to some antibiotics, we quantified the evolution and spread of drug-resistant *E. coli* through spacetime using image analysis and quantitative PCR. As is done in hospitals routinely, we exposed the bacterium to a gradient of antibiotic in a ‘disk diffusion’ drug susceptibility test that we videoed. The videos show complex spatio-genomic patterns redolent of, yet more complex than, Fisher’s predictions whereby a decelerating wave front of advantageous genes colonises towards the antibiotic source, forming bullseye patterns en route and leaving a wave back of bacterial sub-populations expressing AcrB at decreasing levels away from the drug source. qPCR data show that *E. coli* sited at rapidly-adapting spatial hotspots gain 2 additional copies of *acr*, the operon that encodes AcrB, within 24h and imaging data show resistant sub-populations thrive most near the antibiotic source due to non-monotone relationships between inhibition due to antibiotic and distance from the source. In the spirit of Fisher, we provide an explicitly spatial nonlinear diffusion equation that exhibits these properties too. Finally, linear diffusion theory quantifies how the spatial extent of bacterial killing scales with increases in antibiotic dosage, predicting that microbes can survive chemotherapies that have been escalated to 250× the clinical dosage if the antibiotic is diffusion-limited.

## Introduction

Quantifying Fisher’s idea^*1*^ that an advantageous gene moves through space as a diffusive wave is difficult for populations of large organisms. It requires spatiotemporal measurements of a single gene under positive selection, sampled repeatedly from those organisms *in situ*, on an evolutionary timescale with minimal, non-destructive interference from the observer. So, to meet these requirements we turn to single-cell organisms.

We created an observational rig for bacterial populations using a bespoke time-lapse camera and used it to seek Fisher waves in an antibiotic disk diffusion assay that is routinely used to determine the antibiotic susceptibility of bacteria. We videoed a bacterial population expressing a green fluorescent protein (GFP) that had been physically fused to an antibiotic efflux mechanism (protein AcrB) in *Escherichia coli* and by culturing the latter in an antibiotic gradient photographed at regular intervals in appropriate light conditions, we used image analyses, corroborated by qPCR, to determine where high AcrB-expressing antibiotic efflux mutants were at all times. This method quantified an increase in antibiotic resistance that occurred within 24h in this clinical assay. Image data show this was due to a 3-fold increase in efflux pumps per cell appearing within complex spatiotemporal patterns.

An important question for *in vivo* treatments that we sought to quantify using an *in vitro* approach is how does an antibiotic become too spatially diffuse, and so achieve too low a concentration, to inhibit bacteria? Moreover, antibiotics exhibit gradients *in vivo*^*2*^ so how does dose escalation mitigate this effect? Indeed, human infections are treated at high dosages that exceed clinical breakpoints^*3–8*^ yet aggressive chemotherapies can fail,^*9,10*^ whether pathogens have pre-existing resistance or else develop resistance during chemotherapy.^*11–17*^ So, here we take a diffusion-theoretic approach to quantify features of the ubiquitous disk diffusion antibiotic susceptibility assay, paying attention to how very high, super-clinical dosing increases bacterial killing. We revisit this classical problem in microbiology^*18*^ and quantifying bacterial ‘zones of inhibition’ (ZoI) mathematically, thus relating gains in inhibition to dosage increases, providing new quantitative expressions that relate dose to ZoI that are consistent with our imaging data.

## Results

### Part I: Waves and bullseyes from antibiotics: predicting spatial structure from ecological diffusion theory

Intriguingly, patterns are often visible in antibiotic diffusion assays.^*19–21*^ To explain them, we turn to an ecological genetics model written in the spirit of Fisher - the CARS equation: carbohydrate-antibiotic-susceptible-resistant - that captures fundamental features of bacterial growth and antibiotic diffusion:

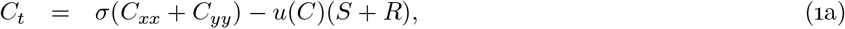

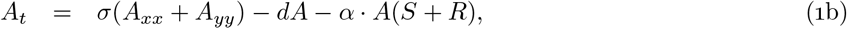

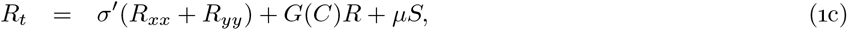

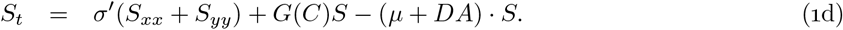

Here, drug-resistant and susceptible bacteria (*R* and *S*, respectively) colonise agar containing a carbohydrate at concentration *C*. This is taken into the cell at rate *u*(*C*) and converted into biomass with efficiency *c*, where all cells have the same half saturation coefficient for resource uptake (*κ*), uptake rate (*u*(*C*)) and maximal uptake rate (*ν*). Thus *G*(*C*) = *c* · *u*(*C*) represents bacterial growth rate, where *u*(*C*) = *νC*/(*κ* + *C*) is carbohydrate uptake rate and *σ* and *σ*′ are diffusion coefficients of small molecules and bacteria, respectively. *μ* is a gain of resistance mutation rate and *D* an antibiotic-dependent death rate and *α* is the antibiotic uptake rate.

In the absence of antibiotic, so that *A* ≡ 0 when *t* = 0, CARS *is* a spatial logistic equation of the type studied by Fisher. To see this, define the number of cells, *N*, by *N* = *S* + *R*. Then define the total system mass as *N* + *cC* and note that the latter is constant through time if *A* = 0, call this constant *C_0_*. Adding (1c) to (1d) yields

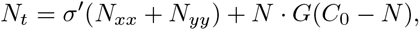

which is a Fisher equation with a vast mathematical history that is known to support waves.^*22*^

#### Bullseye patterns

CARS predicts the formation of spatial patterns when antibiotics are deployed (Figure 1A) because it generates moving bi-modal distributions of bacterial densities formed from waves whereby local population density maxima occur both near to the antibiotic source and far from it, thus creating ‘bullseyes’. Intuition behind the bullseyes is this: high antibiotic dosages kill (or inhibit) so many cells that they create spatial regions abundant in nutrients close to the drug source. So, counterintuitively, the resistant cells that are able to grow in those high-nutrient, low-competition regions can grow more quickly than drug-susceptible cells situated much further from the antibiotic where nutrients are lower per cell because competitor densities are higher there.

**Figure 1:**
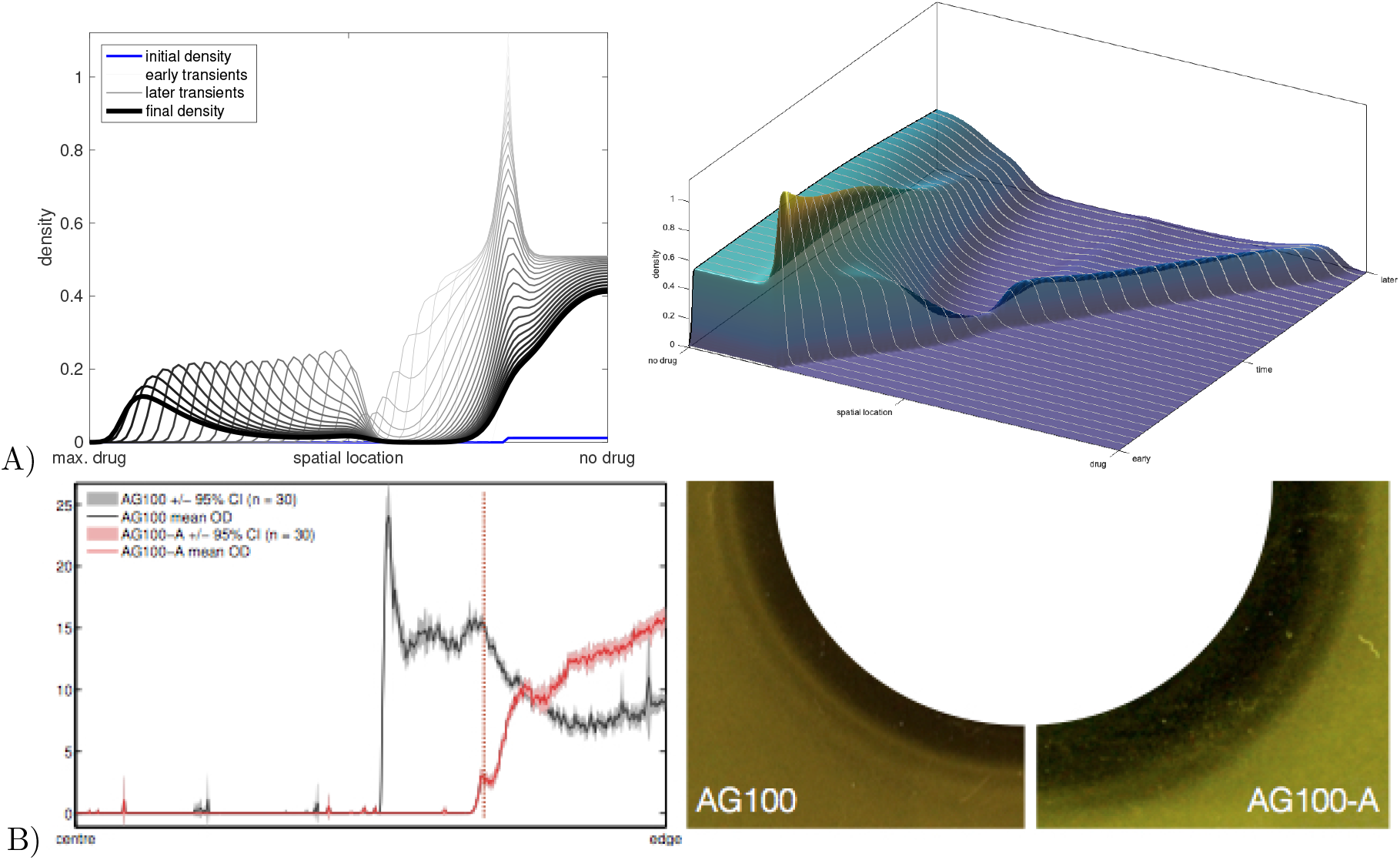
Theoretical predictions: antibiotic diffusion creates bacterial waves and rings. A) Simulating antibiotic growth inhibition on agar using radially symmetric solutions of the CARS equation: a bacterial inoculation localised to the left of the spatial domain where there is no drug eventually forms a bi-modal spatial wave. Note how a sharp population density peak first forms (this creates the first bullseye ring) and then decays while a resistant wave front breaks off from the bulk of cells and propagates into the region of high antibiotic concentration (this creates a second bullseye ring). The left image shows radial slices through simulated cell densities at moments increasing in time, the right image uses those slices to form a surface. B) Bullseyes (meaning local population density maxima) are apparent in empirical imaging data (leftmost) that can be seen by eye (right) in photographs when *E.coli* K12 strain AG100 is challenged by doxycycline; the left-hand data are radial averages of the right-hand assay images. Note the 3 local maxima (i.e. the bullseye rings) in the AG100 density profile. Data for strain AG100-A that does not posses a functional *acr* operon exhibits a less visible ringed pattern in the same experimental conditions with a larger zone of inhibition for the same dose (black region, far right).

This intuition (formalised in Supplementary §1) is borne out by imaging data extracted from photographs of disk diffusion assays (Supplementary §2) where *E. coli* K12 (AG100) was exposed to gradients of doxycycline (Figure 1B), creating bullseye patterns of population densities. Our explanation for this is known as competitive release which exacerbates resistance because resistant mutants, namely ones that can survive high antibiotic dosages, are granted access to more nutrients than would have been the case if the drug were not present.

The theory of competitive release predicts (Supplementary §1) that the bullseye can form from different genomes which exhibit their fastest growth rates at different distances from the drug source, thus one genotype could be largely responsible for one bullseye ring. To test this idea, we sought spatiotemporal data on population densities of a genetic mutation and so we observed *E. coli* K12(eTB108) in a disk diffusion assay using doxycycline (Supplementary §2). Strain eTB108 has GFP physically fused to AcrB and it amplifies the efflux operon *acr* that encodes subunits of the multi-drug efflux pump AcrAB-TolC^*23,24*^ under doxycycline stress because this drug is a substrate of the pump. By photographing the diffusion assay through green filters and using quantitative PCR to determine the number of copies of *acr* per chromosome to corroborate the fluorescence images, we sought to track the spatial locations of *acr* amplification mutants across the agar plate.

Bullseye rings of population densities formed during this assay (Figure 2A). A ring was first measured 160 pixels (NB: 13.2 pixels per mm throughout) from the drug source at 8h, which subsequently transits with diminishing wavespeed (Figure 2B) towards the antibiotic. qPCR data show *acr* per genome was amplified within 24h and *acr* copy number correlates positively with distance to the doxycycline source at 24h (Figure 2C; linear regression *p* < 0.002, *F* ≈ 19.6, AICc ≈ 16.2, quadratic regression *p* ≈ 0.0026, *F* ≈ 12.4, AICc ≈ 18.1). However, the spatial distribution of *acr* per genome had changed quantitatively by 48h and a ‘hotspot’ formed whereby *acr* copy number was now maximised some intermediate distance from the drug, as indicated by a quadratic regression being a more likely descriptor of the data than a linear regression (linear regression *p* ≈ 0.05, *F* ≈ 4.7, AICc ≈ 27.0; quadratic regression with unimodal geometry *p* ≈ 0.0012, *F* ≈ 12.5, AICc ≈ 22.1). Thus, later populations harbour mutants with 2 mean copies of *acr* per genome (and 3 copies were observed in 1 biological replicate) whereas earlier populations have fewer operons (c.f. Figure 2C and D).

**Figure 2:**
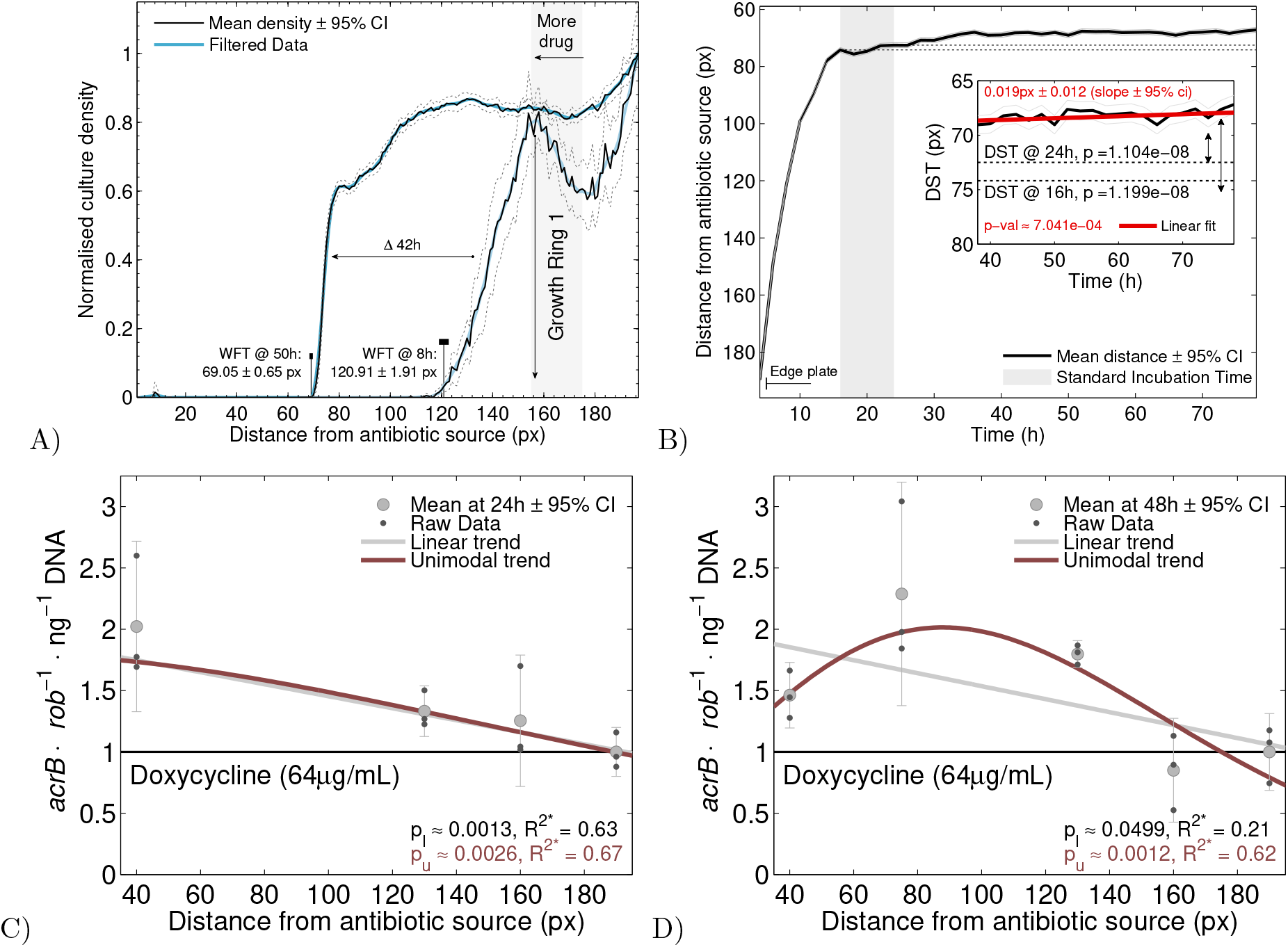
Hotspot dynamics of acr per genome in a doxycycline gradient. A) Spatial dose-response profiles after 8h and 50h of exposure to doxycycline. A bullseye ring was detected at radius 158 ± 3.6 pixels (mean ± 95% CIs, n = 22). The wavefront location (WFT - in pixels), defined as the radius at which 1% of the population density was observed relative to the value at the outer edge of the agar plate, was measured at 121 ± 2 pixels (black line on the x-axis, mean ± 95% confidence, n = 22) after 8h and at 69 ± 0.6 pixels after 50h. B) Dynamics of the front: the inset highlights the position (DST - distance in pixels) of the front at 16, 24h and beyond: a linear regression (shown in red alongside mean slope ± 95% confidence interval) shows the front speed slows in time with a bi-phasic structure whereby the rapid early inroads towards the high-dose region slow after 16h. C) Copy number of *acrB*, a component of the *acr* operon, per chromosome regressed against distance to the drug source (supplied dose as indicated) after 24h. Raw data are black dots, mean and 95% confidence interval are grey. To test for local maxima, *acrB* per cell was modelled using linear and quadratic regressions; *p* and adjusted *R*^2^ values are shown and the linear regression is the more likely datafit here. D) Copy number of *acrB* per chromosome as a function of distance to the drug source after 48h: the quadratic regression is now the more likely datafit. This is consistent with theory (Supplement §1, Figure S1) showing that growth rates of different mutants are maximised at different intermediate spatial locations.

#### Genomic waves

Seeking empirical evidence of waves, we examined relative fluorescence image data (green light intensity per pixel divided by white light per pixel) as a proxy for the spatial distribution of the per-cell abundance of AcrB (Figure 3A shows a positive nonlinear correlation). These data indicate (Figure 3B) an expanding population front at the edge of the ZoI situated approximately 80 pixels from the drug source at 24h where qPCR and GFP imaging data both indicate between 2 and 3 copies of *acr* per genome. Imaging data indicate a spatial gradient of up to 3 AcrB relative protein units per cell where cells have 1 AcrB relative unit per cell at those spatial locations sited as far as possible from the drug source (Figure 3B and C).

**Figure 3:**
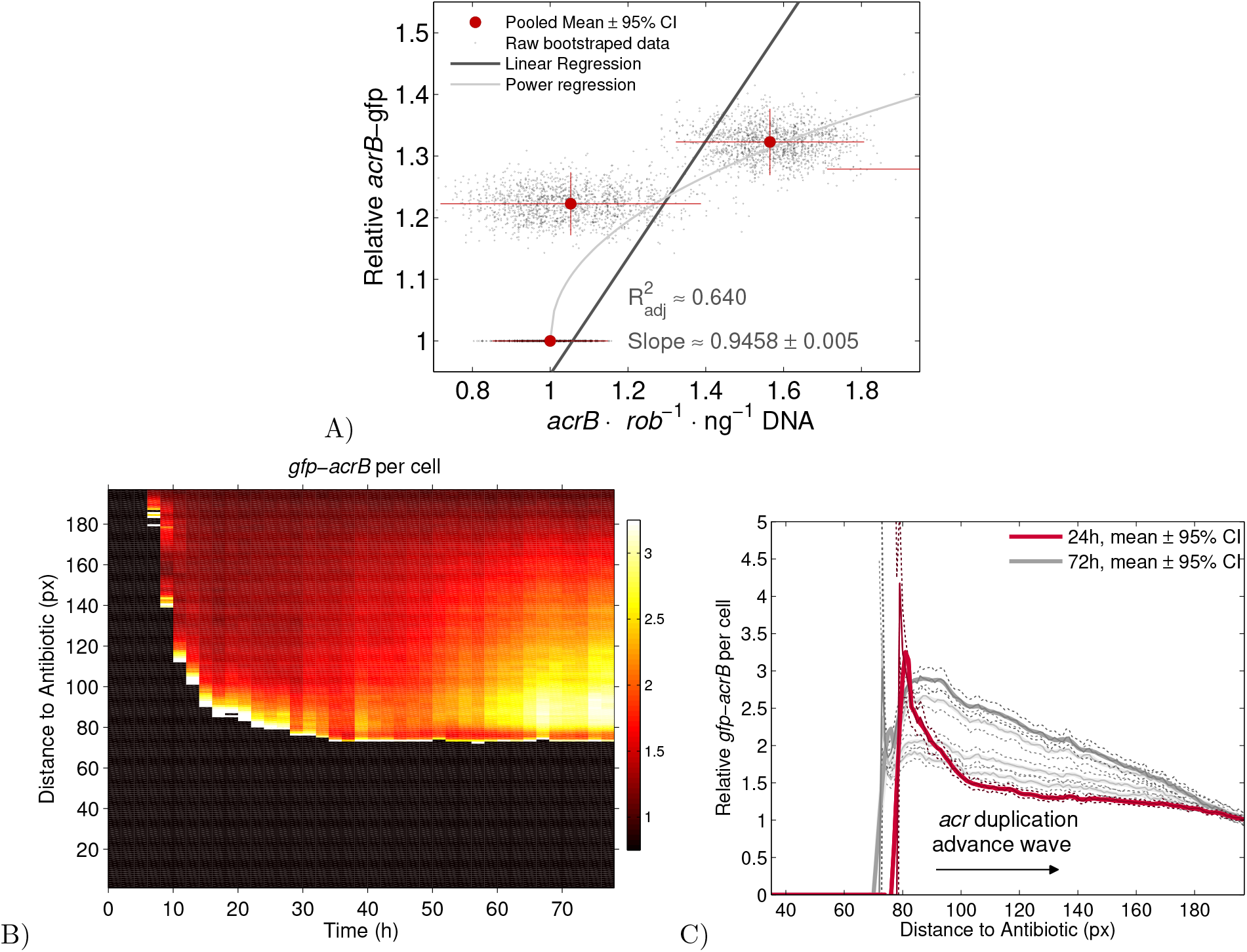
A wave front and back in a doxycycline gradient. A) A significantly positive correlation is observed between the green per white light signal (a proxy for AcrB per cell) and acr operons per genome: y-axis shows green light per white light intensities, x-axis shows acrB per genome from qPCR. Linear and nonlinear (power law) regressions illustrate the positive correlation between these two sets of empirical observations. B) Abundance of GFP-AcrB per cell in spacetime: dark red represents the wild-type, background abundance found at the edge of the plate (unity on the colour gradient) while white represents the highest observed abundance; black represents no growth. C) The distribution of AcrB per cell (green light per white light) between 24h (red) and 72h (dark gray) with data at other times indicated as thin lines: the leftmost peak at 24h subsequently widens whilst AcrB copy number progressively increases, forming a more highly-resistant wave back behind a slowly advancing wave front at later times.

The wave front exhibits a bi-phasic wave speed (Figure 2B) that approaches the antibiotic source at two different rates that slow but do not stop (Figure 2B, linear regression: 0.019 pixels per h ± 0.012, slope ± 95% confidence interval, *p* ≪ 0.001). The faster initial phase (before 16h) is likely driven by the carbon gradient-mediated growth of drug sensitive, wild-type cells with 1 *acr* per genome. After 24h, the emergence of higher *acr* copy-number variants likely forms the second, slower phase of colonisation towards the drug source as mutants overcome both the increase in antibiotic dose at those locations and the large fitness costs of carrying *acr* amplifications.^*24*^ Finally, we highlight a wave-back behind the front, away from the drug source, where AcrB per cell increases towards the antibiotic source (Figure 3C), up to approximately 3 units relative to 1 for the wild-type.

### Part II: how zone of inhibition size scales with dose

We are now interested in how the size of the zones of bacterial clearance depend on the antibiotic dosage supplied. The above theory and data indicate subtle nonlinear dependancies that we henceforth ignore in the expectation that linear diffusion theory might be sufficient to answer this classical question. Indeed, much can be gleaned from basic dimensional considerations alone. For instance, spatially-extended chemotherapy should follow a law of diminishing returns because as antibiotics diffuse through 3D-space, killing as they go, the ZoI radius should depend on the 1/3 (cube) root of the dose applied: to *double* the ZoI volume, *8 times* more antibiotic must be supplied. This number is larger if the antibiotic degrades as it diffuses, as we now show.

#### Diffusion-kill theory

We now present the simplest possible theory of spatially-extended antibiotic killing that we call *diffusion-kill theory*. Assume a spatially extended bacterial population encounters an antibiotic diffusing isotropically from a point source at rate *σ*. Suppose antibiotic is supplied at the centre of a spatial domain at concentration *A_c_* and that the distribution of antibiotic, *A*, is described by the fundamental solution of the diffusion equation in *n*-dimensions (*n* ≤ 3). This is *A_t_* = *σ*(*A_xx_* + *A_yy_* + *A*_*zz*_), where *x, y* and *z* are spatial coordinates, thus:

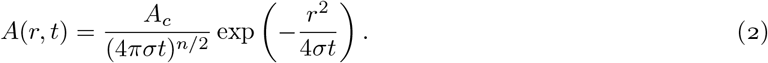

Here, *r* is distance from the antibiotic source so that *r*^2^ is either *x*^2^, *x*^2^ + *y*^2^ or *x*^2^ + *y*^2^ + *z*^2^ for *n* = 1, 2 and 3 respectively and *A*(*r*, *t*) represents a spatially-normal distribution that expands out from the source as a decaying, spherical wave. Experiments correspond to *n* = 2 and *n* = 3 where the former approximates an agar plate where molecules cannot penetrate far into the agar and the latter approximates a region of homogeneous tissue.

Suppose a threshold concentration exists above which the antibiotic kills cells, call this threshold *A_d_* (c.f. the minimal bactericidal concentration (MBC)). The ZoI is described by the set of coordinates at distance *r* from the antibiotic source that satisfy

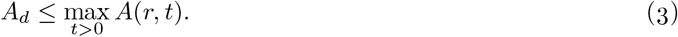

We can solve inequality (3) (Supplement §3) which yields a constant depending on spatial dimension and *A_d_*, call it *C*, whereby killing occurs at those distances, *r*, for which

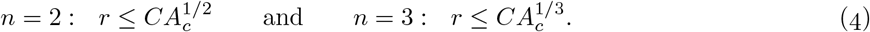

This formulae are expected from dimensional considerations but the calculations generalises to the situation where the drug decays: in 2 dimensions we find constants *C*_1_, *C*_2_ and *C*_3_ (Supplement §3) such that

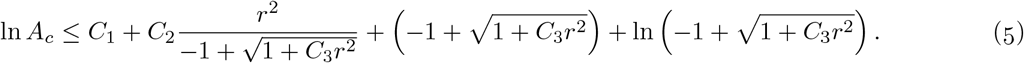

#### Testing diffusion-kill theory

We tested (5) by imaging disk diffusion assays, as follows. We excised a circular section of agar from a plate, replacing it with agar supplemented with identical nutrients but with added antibiotic at a defined dose (Methods). We photographed the latter (Figure S3) to track bacterial inhibition, taking photographs at regular intervals to produce data on the dynamics of the sizes of ZoIs.

The resulting data agree with theory: ZoIs at up to 256×MIC follow the diminishing returns law (4) although the model (5) better fits data for penicillin (Figures 4A, S6) though not for doxycycline (Figures 4B). Prior theory (*25*, equation 1),^*18*^ predicts ZoI geometries of the form *r* ≤ *C*_1_ ln(*A_c_*/*C*_2_))^1/2^ which capture data well too, though not as well as diffusion kill theory (Figures 4A and B; more replicates in Supplement §3). However, prior theory makes a non-physical prediction that the ZoI boundary (where *r* = *C*_1_ ln(*A_c_*/*C*_2_))^1/2^) has a logarithmic singularity that grows without bound as the antibiotic concentration reduces. Our theory corrects this: exponentiating (5) predicts *r* → 0 as *A_c_* → 0, meaning the ZoI disappears when there is no drug.

**Figure 4:**
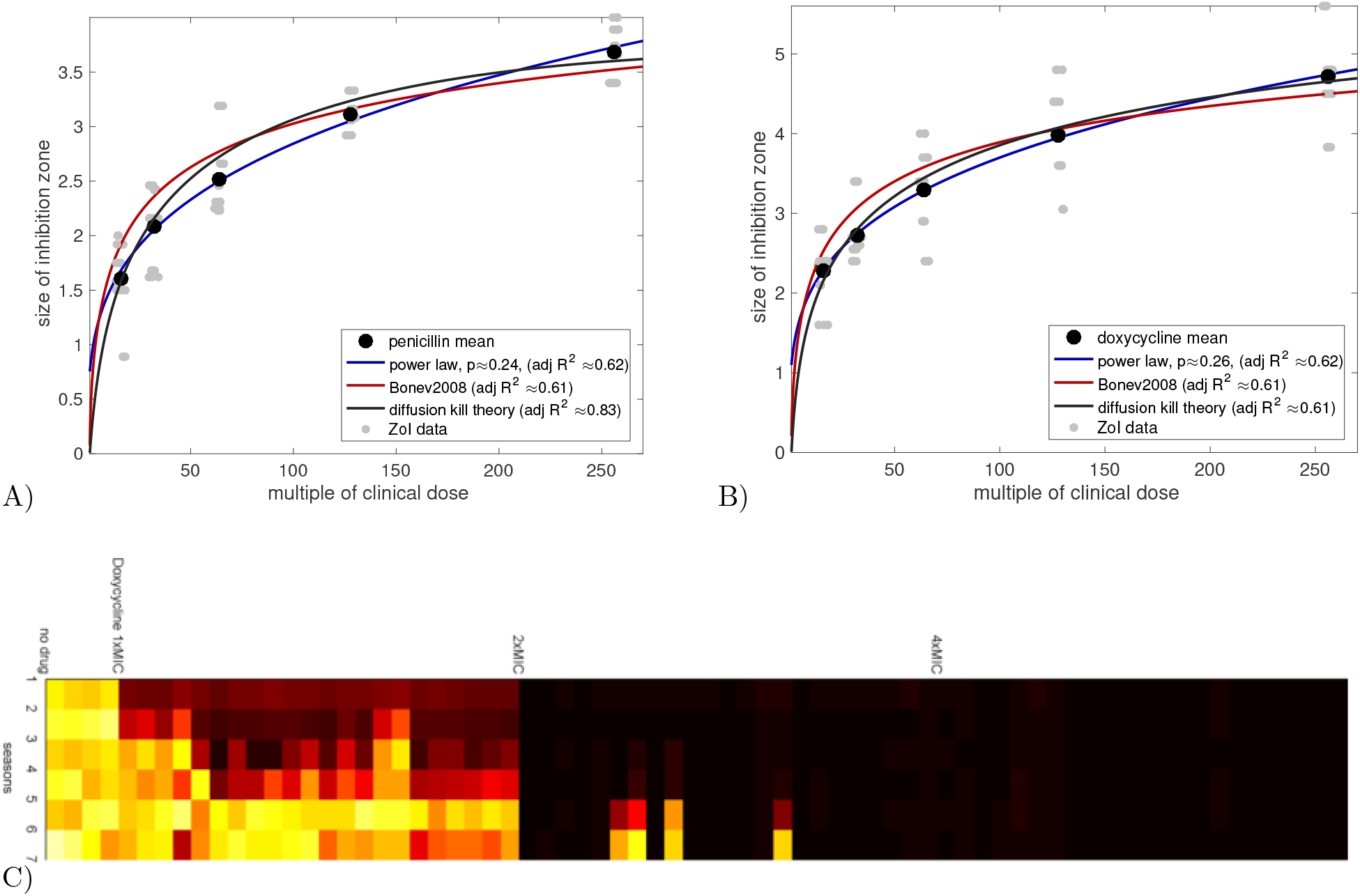
Diffusion theories capture bacterial ZoI data. A) ZoI radii regressed against antibiotic dose data using penicillin and *E. coli* K12 (MG1655). The regression using (5) is called ‘diffusion kill’ in the legend, the model from (25) is called ‘Bonev2008’ and a power law regression is motivated by (4). Adjusted *R*^2^ values show all models provide similar quality datafits. B) Is the analogy of A but for doxycycline where (5) provides the most appropriate fit to data. C) A heatmap showing the population density (optical density) of *E.coli* K12 (MG1655) propagated in liquid media at increasing dosages of doxycycline. Integer multiples of the MIC used during 7, 24h-long growth ‘seasons’ are indicated where data to the right of the MIC marker were obtained in that antibiotic background. Population density was measured at 24h each treatment round and the hot colour scale indicates white/yellow as high and red/black as low, columns denote different replicates and rows denote 24h seasons. Note when dosage is 4×MIC we observe no growth (black regions), although 1×MIC and 2×MIC support increasing growth.

## Discussion

Theory and data highlight a fundamentally different bacterial dose response to antibiotics in spatial assays compared with non-spatial assays conducted in mixed liquid environments. Notwithstanding the Eagle effect^*26*^ whereby dose-response data can be non-monotone in liquid cultures, methods for determining antibiotic dose responses generally predict that bacterial growth declines with increasing antibiotic dose. Here, in contrast, spatial structures form when antibiotics diffuse from a source so that bacteria can grow fastest in a mid-dose regime, leading to patterned growth within waves and rings of colonisation moving towards the antibiotic source at slowing rates.

One mechanism supporting non-monotone spatial dose responses is competitive release: antibiotic gradients create nutrient gradients, thus cells that can grow in zones rich in antibiotic (i.e. resistant cells) access more nutrients than cells far from the antibiotic source. This creates conditions whereby the most rapid population growth can arise close to (and not far from) the antibiotic source because growth benefits provided by the nutrients there outweigh any costs of being resistant. Here, this dynamic promotes resistance by producing cells with 2 additional copies of the *acr* efflux operon within 24h in a mid-dose region. We tracked these mutants in real-time by combining qPCR with image analysis of GFP-labelled AcrB protein and observed gradients of *acr* amplifications in a wave structure that qualitatively resembles Fisher’s predictions,^*1*^ albeit with additional complexities, like biphasic wave speeds and bullseye rings.

Antibiotics form gradients in the body ^*2*^ so predictions on resistance progression *in vitro* can only have limited predictive power *in vivo*. Our data is consistent with this idea in the sense that spatial and non-spatial assays behave differently: here, the evolution to 3-fold *acr* amplifications observed within 24h in a spatial antibiotic gradient dosed at 128× MIC compares unfavourably against the mere 4×MIC needed in a shaken liquid environment to prevent detectable bacterial growth, therefore preventing *acr* amplifications, following 7 rounds of antibiotic treatment (Figure 4C).

In conclusion, fluorescence photography and imaging data extracted from a ubiquitous clinical assay show a 250-fold increase in antibiotic concentration increases bacterial clearance by much less than 1cm on an agar plate. Thus, without active transport mechanisms to a bacterial site or else spatial mixing to prevent gradients forming, diffusion can prevent antibiotics from achieving the high concentrations needed to kill bacteria. Drug diffusion can therefore promote resistance by providing a smooth path that leads from the lowest to the highest dosages over short spatial scales.

## Methods

### Media and strains

We used strains *E. coli* K-12 MG1655 and TB108. Strain TB108 derives from strain MG1655 in which a superfolder GFP locus was inserted in-frame and downstream of *acrB*. As a consequence, the GFP is attached to the cytoplasmatic c-terminal end of AcrB after translation, with an additional eight amino acids long polylinker peptide between AcrB and GFP. The fluorescence tag led to a partial malfunction of the pump AcrAB-TolC, as determined by the MIC of the strain TB108 to erythromycin, that was repaired (increased back to wild-type) by propagating TB108 in M9 media supplemented with 0.2% glucose (w/v), 0.1% casamino acids and 10*μ*g/mL of erythromycin for seven days. After this step, the MIC of the evolved TB108 strain (now denoted eTB108) was identical to the wild-type MG1655. By using erythromycin we sought to prevent mutations that pre-adapt eTB108, that we use used to the track abundance of GFP-AcrB, to doxycycline and penicillin. The same M9 media, but with no antibiotic, was used for Petri dish assays by adding 6g/L of agar powder (semi-solid agar). To embed the microbes in semi-solid media, 1mL of overnight culture in liquid media was added to 99m L of semi-solid media prior to use.

### Spatially-extended diffusion assays

For spatial diffusion plates we prepared two sets of semi-solid media, one being inoculated from an overnight culture as described above, and the other containing doxycycline equivalent to from 1 to 300 times the minimum inhibitory concentration (1×MIC = 0.227 ± 0.007*μ*g/m L, mean ± s.e.m. with n = 8). To our theoretical assumptions, a volume of 20mL from inoculated semi-solid media was used to fill the Petri dish and we later removed a circular section, at the centre of the Petri dish, we then refilled this with semi-solid media containing the drug (1.3m L used). The resulting concentration of nutrient was therefore uniform at the start of this assay whereas that for the drug was not because the drug source lay at the centre of the plate.

### Quantitative polymerase chain reaction (qPCR)

We first removed a section of agar of 2.5×5×5mm, approximately, at different distances from the source of antibiotic and degraded the agar using agarase (ThermoScientific #EO0461) as per manufacturer instructions but with additional incubation time. We then extracted DNA from each fragment using the DNA extraction kit ‘GeneJET’ (ThermoScientific #K0729), quantified DNA yield using ‘Qubit’ and diluted accordingly to harmonise DNA content. The kit ‘Luminaris Color Probe Low ROX’ (ThermoScientific #0342) was used as per manufacturer instructions to quantify the abundance of the loci *acrB* and *rob*, the latter being a housekeeping gene of reference, sited outside the genomic region known to be amplified under our experimental conditions. The amplification efficiency was ≈ 100% with the primers and probes described in Table 1 and, based on the resulting calibration curves (Figure S7), we used 25ng of material from the agar samples.

**Table 1:**
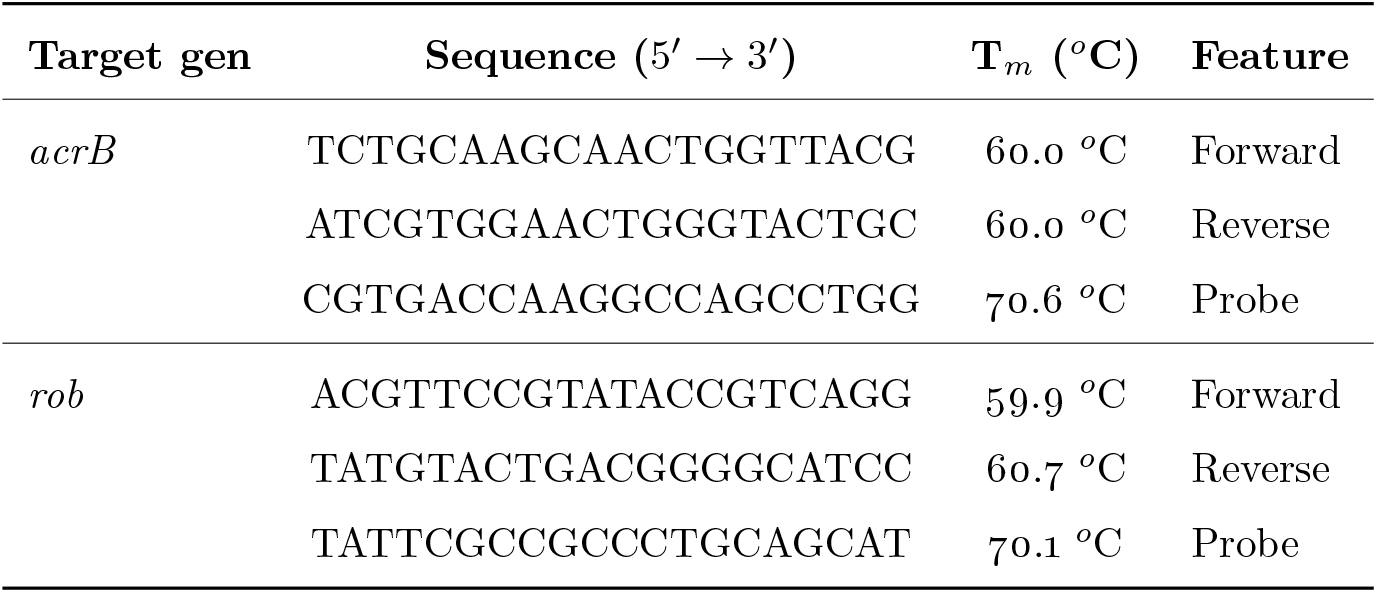
Primer and probe sequences for the loci *acrB* and *rob*. The amplicon size for *acrB* is 104bp and 112bp for *rob*.

### Image analysis pipeline

To track culture growth we used a custom-built plate imaging system (Figure S3) equipped with a digital camera (Canon EOS 1100D). To detect GFP-AcrB, we used high intensity LEDs with an effective excitation bandwith of 475-490nm (Comar #475GY25 and #490IK25) and filtered emission bandwiths in the 530-560nm range (Comar #530GY50, #560IK50). We implemented an algorithm in MATLAB to read the resulting time lapse data, with photographs taken every 2h for three days, thus determining dynamic, spatial dose-responses. After a baseline correction to harmonise readings from subsequent photographs, we generated a dose-response profile measuring the pixel intensity in a 16-bit scale (0-255 for each red, green and blue channel) across a line connecting the centre of the zone of inhibition to the edge of the Petri dish, and rotated this line every 15° to generate technical replicates.

### Data analysis

To measure the effect of drug dose on microbial growth, we fitted m athem atical models to population density and growth data to determine dose-response profiles (Figure S4A). The following logistic model was used to model bacterial density timeseries at different distances from the drug source:

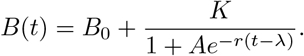

Here *B*_0_ is an estimated experimental blank due to reading an empty microtitre plate, *K* is the culture density during stationary phase or *carrying capacity, A* is a composite parameter that includes the initial cell density (inoculum), *r* the growth rate per cell per hour and λ the duration of the lag phase in hours. Curves for GFP-AcrB were reconstructed (Figure S4B) by fitting the following model to timeseries data

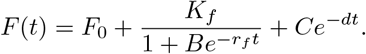

The constant and logistic terms here are analogous to those in the above population growth model and the additional exponential term *Ce*^−*dt*^ models the potential for down-regulation or degradation of GFP-AcrB, at rate *d*.

## 1 Supplementary Information: Growth rate is maximal an intermediate distance from an antibiotic source

One can use the competitive release concept to explain bullseye patterns quite directly. To do so, modify the basic Monod growth law from the main text and represent bacterial growth rate (*G*) as

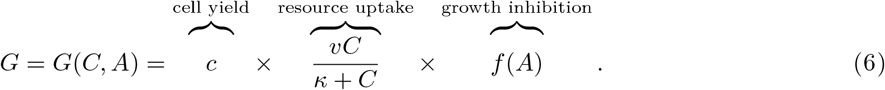

where *G* is reduced by a function of the antiMicrobiol, *A*, by an inhibition function, *f*, that satisfies 0 ≤ *f*(*A*) ≤ 1. Assuming *A* and *C* diffuse, we can predict how *G* changes through space assuming two things: the drug decays and carbohydrates create the conditions that exhibit competitive release at very high drug dosages. For this, write the steady-state solution of the decay-diffusion equation *A*_*t*_ = *σ*(*A*_*xx*_ + *A*_*yy*_) − *dA*, as *A*(*r*) = *A*_*c*_*ϕ*(*r*), where *d* is the antibiotic degradation rate and *A*_*c*_ is the antibiotic supply concentration. Assume the free diffusion of carbohydrate away from the antibiotic source occurs due to the use of high antibiotic dosages, so the steady solution for *C* is analogous: *C*(*r*) = *C*_0_*ϕ*(*r*), where *C*_0_ is the carbon supplied that is initially uniformly distributed on agar at *t* = 0.

If one could show these assumptions imply the maximal bacterial growth rate occurs at intermediate spatial locations sited away from the antibiotic source and if the intracellular antibiotic concentration for two sub-populations were different, for instance if one were to express more antibiotic efflux pumps than another, then their spatial location of optimal growth would be different too. Thus bullseyes would form from the growth rings of the different sub-populations.

So, to demonstrate this, take growth rate defined in (6) and suppose that the spatial distribution of *A* is given by the steady-state solution of the diffusion equation,

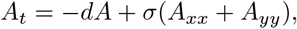

so *A* = *A*_*c*_*ϕ*(*r*) where *ϕ*(*r*) is a strictly decreasing function. So *ϕ*(*r*) satisfies, for this example,

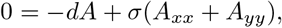

(although our argument cam be made much more general than this single model). Now assume that where the drug source is, doses to cells are so high so they are rapidly killed due to the high dose, thus a carbon ‘haven’ is formed which begins to supply carbon to surrounding cells by diffusion. Thus, *C* is also the steady-solution of a diffusion equation and *C* = *C*_0_*ϕ*(*r*). In the following, a dash (′) and a subscript (typically *r*) will both denote derivatives, as is standard notation.

Growth rate is maximal where *∂G*/*∂r* = 0 and ln*G* = ln *v* + ln (*f*(*A*)) + ln (*C*) − ln (*κ* + *C*), so that

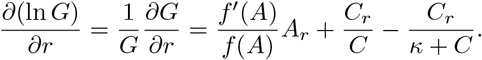

Using *f*(*A*) = 1/(1 + *pA*) for the sake of definiteness, we find *f*′(*A*) = −*p*/(1 + *pA*)^2^ so that *f*′(*A*)/*f*(*A*) = −*p*/(1 + *pA*) and therefore

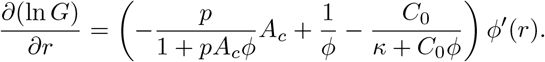

The assumed monotonic property of *ϕ* (assuming the drugs decay away from the source, which is consistent with solutions of diffusion equations) now means that we are seeking the values of *r* for which

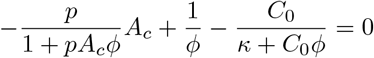

where *ϕ* = *ϕ*(*r*). Simplifying this, we arrive at the following fixed point equation for *ϕ* = *ϕ*(*r*):

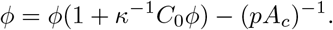

Elementary sign and growth considerations of the RHS of this equation (which is quadratic in *ϕ*) show it always has a unique solution, *ϕ*\*, so that the point of maximal growth rate in the spatial domain is the value *r* = *r*\* for which *ϕ*(*r*) = *ϕ*\*.

Now note that if a population of cells has two different responses to antibiotics whereby each cell is associated with a different internal drug concentration, then this is represented in this model by having *A* take on lower values for the more resistant cells. For instance, having more drug efflux pumps would reduce *A* when is modelled here by scaling the value of *A*_*c*_ by a constant between 0 and 1, call this λ: *A* = λ*A*_*c*_*ϕ*(*r*) is the internal concentration of antibiotic of a cell at location *r* where *λ* is lower for cells with more efflux pumps. Thus the equation for maximal growth rate becomes

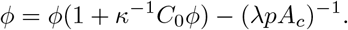

Now, as the resistance phenotype becomes increasingly significant, represented by *λ* reducing towards 0, the form of these equations makes the value of *ϕ*\* become larger and so *r*\* becomes inreasingly smaller because *ϕ*(*r*) is a decreasing function. It is feasible that the value of *ϕ*\* becomes so large that *r*\* approaches 0 and so the fastest growth rate of some highly resistant strains could be achieved very close to the drug source.

Continuing this argument, because of the efflux from highly resistance cells, it could arise that there is a local increase in drug concentration within some highly drug-susceptible cells above the value *A*_*c*_. In this case, *λ* would be bigger than 1 and, as this value grows, the solution *ϕ*\* heads towards zero and so *r*\* could grow towards being a point very far from the drug source.

To get a sense of how the resulting spatial distributions of population density might appear, consider the mathematical solutions provided by the above analysis. So, first we need solutions o f the radially symmetric diffusion equation where *x* and *y* coordinates are replaced by the distance, *r*, from the drug source and the angular coordinate is removed: 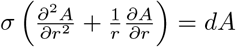, such that *A*(*r*) → 0 as *r* → ∞ and *A*(*r* = 0) = *A*_*c*_, or *A*_*rr*_ + *r*^−1^ *A*_*r*_ −*Ad*/*σ* = 0. As the solution of this equation is a Bessel function that requires lengthy techniques to derive that are likely to obscure our point, we illustrate the 1-dimensional version ofthiscalculation in Figure S1 where *A*(*r*) is an exponential function *A*(*r*) = *A*_*c*_ exp(−*r*(*d*/*σ*)^1/2^).

**Figure S1:**
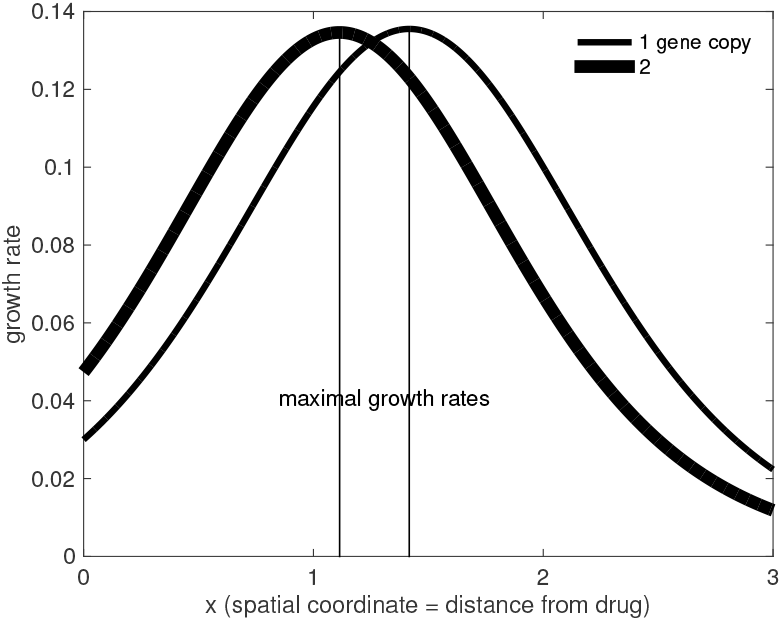
Two different theoretical growth rate profiles for bacteria carrying different numbers of a drug resistance gene that carries costs based on the Monod-like analysis in this section. Note how the addition of the gene shifts the growth rate maximum point towards the drug source but makes growth slower further away from the source.

When we compare the theoretical Figure S1 against the empirical equivalent of the same concept, we find Figure S2 which has been determined for the total population, not for a sub-population. Thus growth rate is maximised some intermediate distance from the drug in data, as the theory claims.

**Figure S2:**
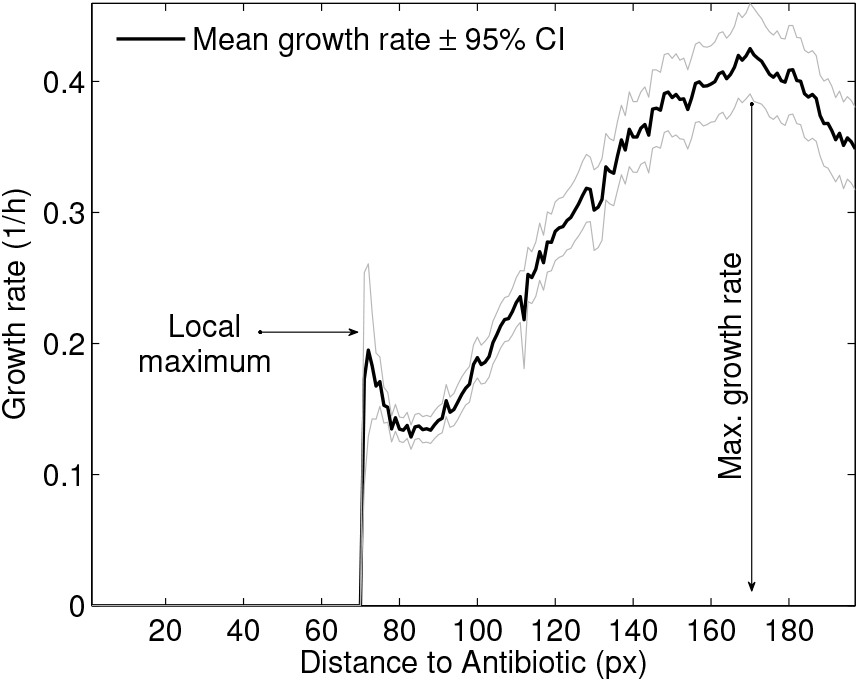
Empirical *per capita* growth rate at different distances from the drug source. The absolute maximum growth rate was detected at 168 ± 2.66 pixels from the source (mean ± 95% confidence interval) and a local maximum is found at 72 ± 0.61 pixels (see Figure S1 for the theoretical derivation of this property).

## 2 Supplementary Information: a fluorescence and white-light photography rig

We constructed a fluorescence photography rig to take images and videos in laboratory conditions. This is based on a standard camera, LEDs used to excite bacterial samples on agar plates in white light and at specific wavelengths, and light filters that are rotated into place by a servo to read different protein emission wavelengths. A heating mat is placed below the camera on a mounting point to control agar temperature.

Antibiotic disk diffusion assays are illustrated in Figure S3B which shows a circular piece of agar (in black) into which high dosage antibiotic has been impregnated which diffuses outwards to create a zone of inhibition on a bacterial lawn. We can measure this lawn either in white light to determine population density, or else in a fluorescent wavelength if wish to determine the level of expression of a GFP-tagged gene throughout space by that population. Dividing white light measurements by GFP measurements gives a proxy for local expression levels per cell through space. Figure S4 shows two exemplars of these spatiotemporal datasets.

### NB

Figure S5 shows that there is a positive, albeit nonlinear and saturating correlation between plated cell counts and white light intensity as measured using a camera. Thus white light intensity can be used as a proxy for cell densities provided the latter is not too high.

**Figure S3:**
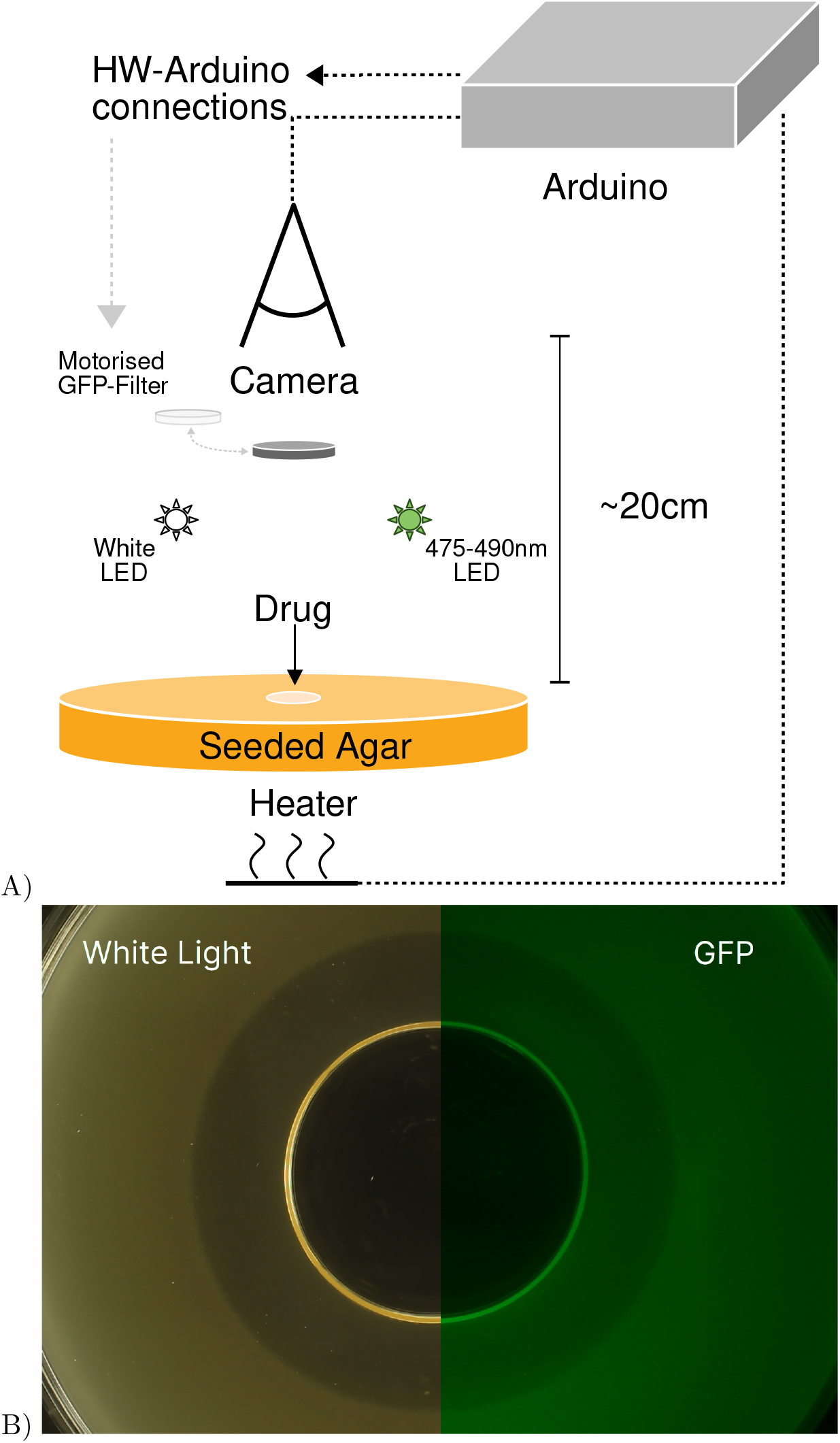
Imaging device. A) Overview of the imaging rig with the control unit containing Arduino UNO on top, from which all the connections stem. Seeded agar plates are placed by a human operator at the centre of the device. B) Sample photographs of a seeded agar plate are used to determine bacterial density (labelled ‘White Light’) and green fluorescence emitted by green fluorescence protein (GFP).

**Figure S4:**
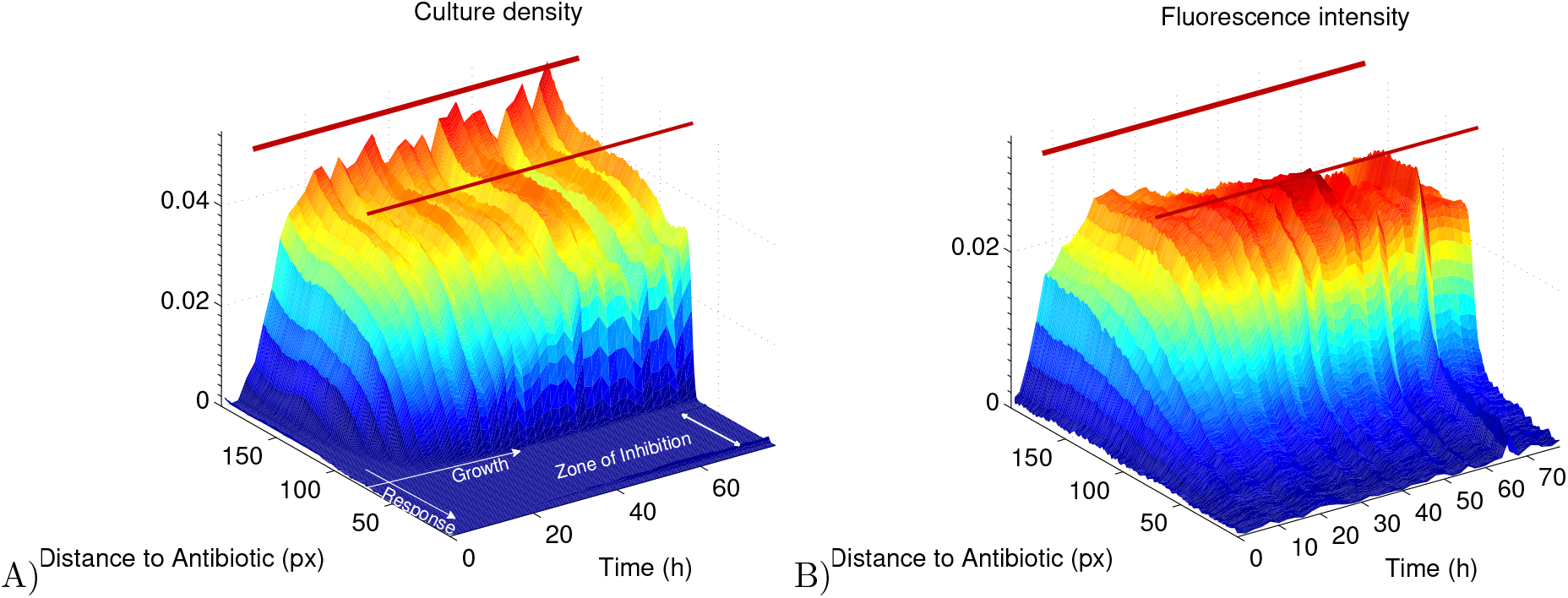
Spatiotemporal density and AcrB∷GFP expression surfaces from image analysis. A) An aggregation of radially averaged white light dose-response data needed to reconstruct one set of technical replicates of growth curves and dose responses, all laid together to form a surface. A lack of growth is shown in deep blue whereas the maximal growth is shown in red. The parallel red lines remark the coordinates at which the absolute and local maximum growth rate were observed for this data whereby *E.coli* K12 (eTB108) is incubated for over 60h at 37°C. B) Analogous data to A) but instead of using population growth data it uses GFP that is physically fused to AcrB (see Methods).

**Figure S5:**
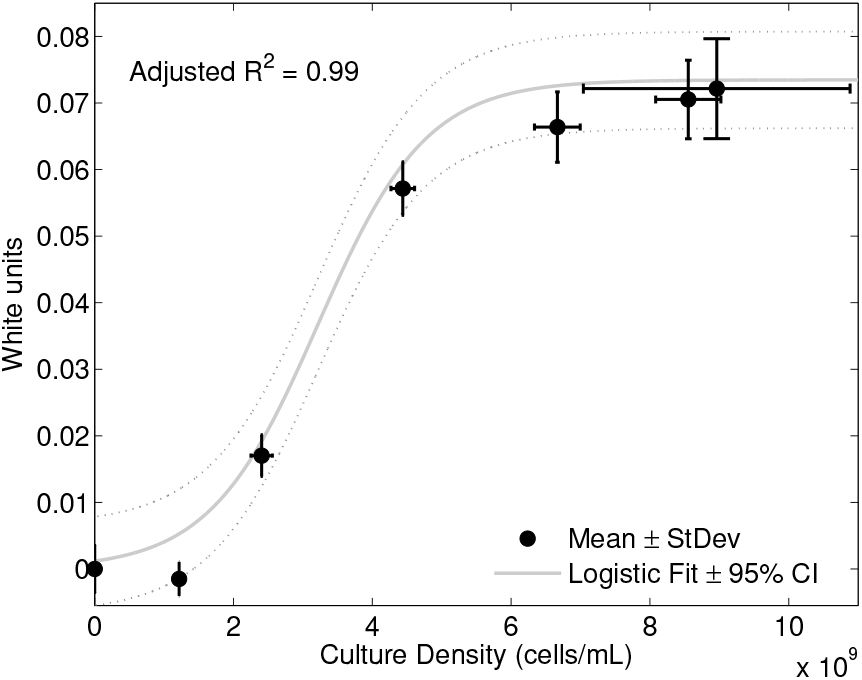
Correlation between white light turbidity and culture density (colony forming units (CFUs)). White light measurements from a standard camera quantified using JPEG RGB values (y-axis) and culture density as CFUs on the same agar plate (x-axis); note the positive, nonlinear correlation.

## 3 Supplementary Information: How zone of inhibition size scales with antibiotic dose

In this section, we use inequality (3) to predict the effect of dose escalation on the size of the zone of growth inhibition. By calculus, differentiate the expression for *A*(*r*, *t*) from the main text with respect to *t*, set this derivative to zero and solve it for *t*. This shows that the maximal drug dose is reached when time, *t*, satisfies *t* = *t*\* = 2*r*^2^/(4*nσ*), at this time 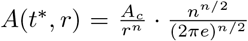. Placing this expression in (3), antibiotics inhibit growth provided 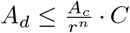, where *C* denotes a dimension-dependent constant. Setting *n* = 1, 2 and *n* = 3 and re-arranging, respectively, we determine the radii of the zones of inhibition: microbes do not grow if they are placed at distances, *r*, from the drug source if

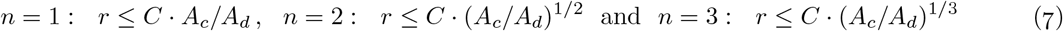

where *C* is a placeholder for constants of proportionality.

We now generalise this calculation to the more realistic case where the drug decays. We again need solutions of the inequality (3) but now where *A*(*t*, *r*) is a solution of the isotropic decay-diffusion equation which, with decay rate *d*, in 3-d reads:

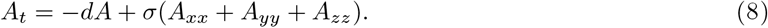

The solution of this is the solution (2) from the main text multiplied by a decaying exponential, at rate *d*:

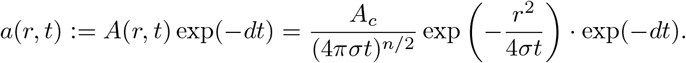

By calculus, we differentiate *a*(*r*, *t*) with respect to *t*, set this derivative to zero and solve it for *t*:

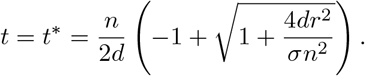

Thus, *a*(*t*, *r*) > *A*_*d*_ for some *t* > 0 if ln *a*(*t*\*, *r*) > ln *A_d_*, or

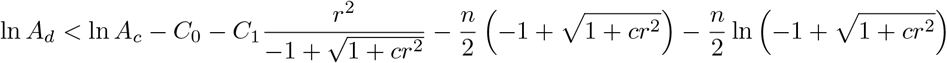

where *c* = 4*d*/*σn*^2^, *C*_1_ = *d*/2*σn*, *C*_0_ = *n* ln(2*πσn*/*d*)/2. Given zone of inhibition radius data on from agar plate diffusion assays, where *n* = 2, this provides a 3-parameter nonlinear regression model to fit against data of the form (re-stating (5) from the main text)

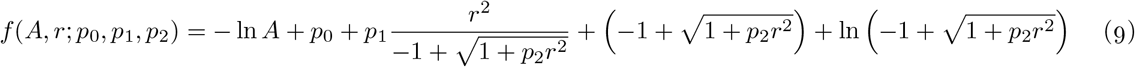

where *A* is the antibiotic supply concentration, *r* is the zone of inhibition radius and *p*_1_, *p*_2_ and *p*_3_ are unknown parameters to be determined from data.

Dimensional arguments from main text suggests the use of the 2-parameter regression model

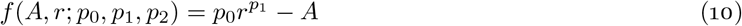

for the same ZoI data where we expect *p*_1_ to be close to the value 2 for diffusion assays on agar plates. This may not be true in practise because agar plates are not 2-d objects, but anisotropic, 3-d objects which are much wider than they are high.

We tested the ZoI regression models defined by equations (4), (9), (10) and one used already in the literature^*18,25,27*^ defined as

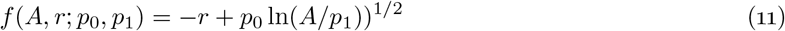

against ZoI data. The result of applying these 4 regressions can be seen in Figure S6.

### NB

Throughout, the equation *f*(*A*, *r*; **p**) = 0 determines the ZoI boundary as a function of dose (*A*) and this observation is used to determined the resulting model parameters **p**.

**Figure S6:**
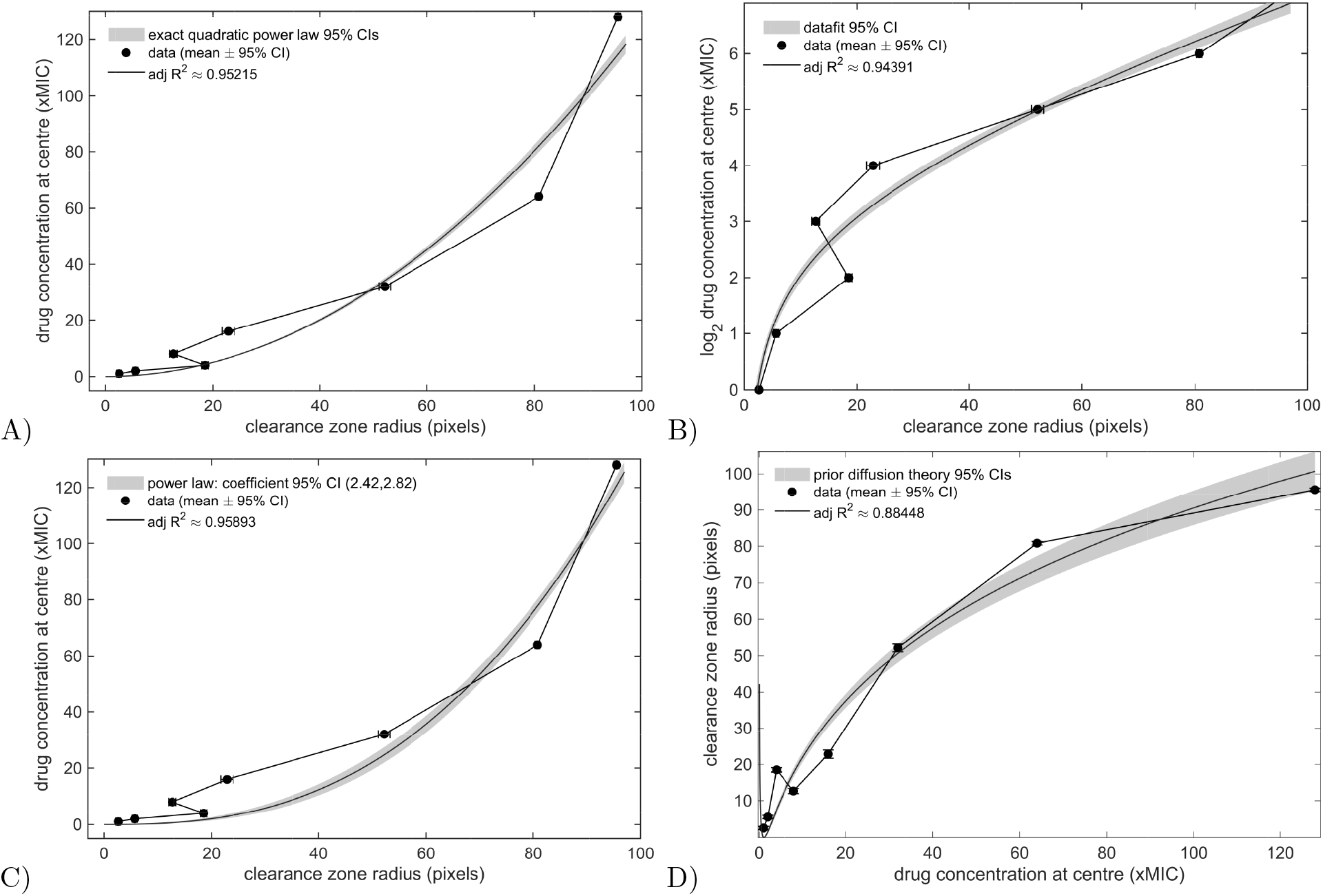
A, B, C and D are the results of applying the regressions (4 - using dimension *n* = 2), (9), (10) and (11) respectively to the penicillin disk diffusion test (as per Figure 4A) repeated for K12 strain AG100. Each plot looks different because the x- and y-axis were chosen to reflect the nature of each regression, some of which use logarithmically transformed data and some which do not. In A, ZoI radii for doxycycline AG100 follow the predicted quadratic power law between radius (x-axis) and dose (y-axis); this is consistent with equation (4) (**adj***R*^2^ ≈ 0.95, F-statistic versus constant model is *F* > 10^3^, *p* ≈ 10^−76^). In all cases, theory and data correlate well but the diffusion-kill theory (9) presented in the main text outperforms prior theory, as demonstrated by adjusted *R*^2^ values in the legends.

Figure S6 shows that the four different possible regressions for the same phenomenon that we propose all capture data, having adjusted *R*^2^ values above 0.88. However, while equation (9) predicts the ZoI radius at the MIC dose, equation (11) makes an unusual and non-physical prediction whereby as dosage heads towards zero, the radius of the ZoI diverges to infinity (see Figure S6). This paradoxical behaviour (of very good fitting at high dose and non-physical behaviour at low dose) can be seen in Figure S6D.

## 4 Supplementary Methods

Methodological data for determining quantitative PCR amplification efficiency are shown in Figure S7.

**Figure S7:**
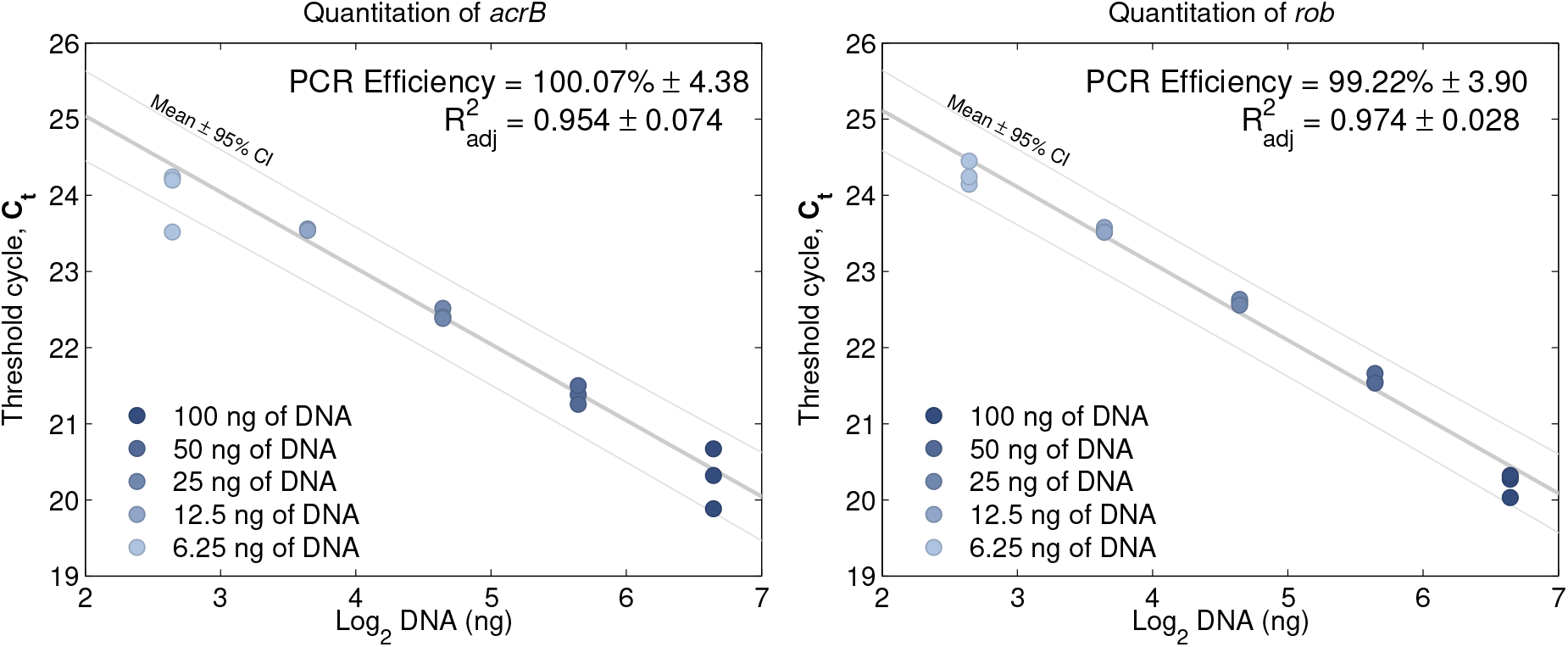
Quantitative PCR amplification efficiency. Calibration curves for *acrB* (left) and *rob* (right). The threshold cycle (*C*_*t*_) is shown on the *y*–axis as a function of DNA content on a log_2_ scale. A linear model was robustly fitted to data (in grey; mean prediction ± 95% confidence interval, *n* = 3). The amplification efficiency was calculated as 2^1/Slope^ − 1, where the slope is given by the linear fit.

